# Reversing gut microbiome-driven adipose tissue inflammation alleviates metabolic syndrome

**DOI:** 10.1101/2022.10.28.514267

**Authors:** N. K. Newman, Y. Zhang, J. Padiadpu, C. L. Miranda, A. A. Magana, C.P. Wong, K. A. Hioki, J. W. Pederson, Z. Li, M. Gurung, A. M. Bruce, K Brown, G. Bobe, T. J. Sharpton, N. Shulzhenko, C. S. Maier, J. F. Stevens, A. F. Gombart, A. Morgun

## Abstract

The gut microbiota contributes to macrophage-mediated inflammation in adipose tissue with consumption of an obesogenic diet, thus driving the development of metabolic syndrome. There is a need to identify and develop interventions that abrogate this condition. The hops-derived prenylated flavonoid xanthohumol (XN) and its semi-synthetic derivative tetrahydroxanthohumol (TXN) attenuate high-fat diet-induced obesity, hepatosteatosis and metabolic syndrome in C57Bl/6J mice. This coincides with a decrease in pro-inflammatory gene expression in the gut and adipose tissue, together with alterations in the gut microbiota and bile acid composition. In this study, we integrated and interrogated multi-omics data from different organs with fecal 16S sequences and systemic metabolic phenotypic data using a transkingdom network analysis. By incorporating cell type information from single cell RNA-seq data, we discovered TXN attenuates macrophage inflammatory processes in adipose tissue. TXN treatment also reversed levels of inflammation-inducing microbes, such as *Oscillibacter valericigenes*, that lead to adverse metabolic phenotypes. Furthermore, *in vitro* validation in macrophage cell lines and *in vivo* mouse supplementation showed addition of *O. valericigenes* supernatant induced the expression of metabolic macrophage signature genes that are downregulated by TXN *in vivo*. Our findings establish an important mechanism by which TXN mitigates adverse phenotypic outcomes from diet-induced obesity and metabolic syndrome. It primarily reduces the abundance of pro-inflammatory gut microbes that can otherwise promote macrophage-associated inflammation in adipose tissue.

## Introduction

Metabolic syndrome (MetS) is associated with abdominal obesity, hypertension, dyslipidemia, and impaired glucose tolerance. It is an increasing global epidemic[1] leading to Type 2 diabetes (T2D) and non-alcoholic steatohepatitis (NASH), recently termed metabolic-associated fatty liver disease (MAFLD) [2-4]. Healthy lifestyle changes and regular exercise along with sustained weight management are the only options to prevent or mitigate severity of the associated diseases as there is no FDA approved treatment; therefore, there is an urgent need to discover additional approaches to prevent or treat MetS [1, 5]. Polyphenols are abundant organic compounds found in plants and used by Ayurvedic practitioners and in traditional medicine for thousands of years to promote various health benefits. The public has great interest in the potential for using these compounds to protect against obesity, metabolic and cardiovascular diseases [6-8]. It has previously been shown, both by us and other groups, that both the prenylated flavonoid, xanthohumol (XN), isolated from hops (*Humulus lupulus* L.), and its semi-synthetic derivative tetrahydroxanthohumol (TXN) reduce biomarkers associated with MetS in animal models of diet-induced obesity. Although both XN and TXN successfully reduce inflammation [9-11] obesity, hepatosteatosis, and insulin resistance [12-16], TXN attenuates these symptoms more effectively than XN [16]. Previously we showed TXN treatment decreases expression of many pro-inflammatory cytokines and hypothesized TXN reduces the infiltration of inflammatory macrophages into adipose tissue [17]. Furthermore, previous studies indicated the gut microbiome as a key factor in the beneficial effects of these compounds [17, 18]; however, the causative microbes and the molecular mechanisms by which they affect the host remain unknown. Therefore, we acquired transcriptomic data from the liver, adipose tissue, and ileum, 16S rRNA sequencing of the fecal microbiome, fecal bile acid levels, and systemic metabolic markers from mice fed a low-fat diet (LFD), high fat diet (HFD) or HFD containing XN or TXN from our previous study [16]. We integrated this data using a data-driven systems biology approach. Through reconstruction and interrogation of a transkingdom network, followed by additional *in vivo* and *in vitro* experiments, we established TXN mediates its primary therapeutic effect by repressing obesogenic diet-associated microbes (e.g. *Oscillibacter* spp.), which consequently decreases metabolically harmful macrophage-associated inflammation in adipose tissue.

## Methods

### Animals and diets

Animals and diets were described previously in detail[16]. Briefly, after acclimation, 60 eight-week-old C57BL/6J mice purchased from The Jackson Laboratory (Bar Harbor, ME, USA) were randomly assigned to one of five diet groups (n=12/group): 1) low fat diet (LFD), 2) high fat diet (HFD), 3) HFD + 0.035% XN (LXN), 4) HFD + 0.07% XN (HXN) or 5) HFD + 0.035% TXN (TXN) for 16 weeks. Body weight and food intake were assessed weekly [16]. Upon conclusion of the study, mice were fasted for 6 h during the dark cycle, euthanized and blood and tissues collected as described previously [16]. One mouse from the TXN group died and was excluded from subsequent analysis. For isolation of stromal cells from visceral adipose tissue, seven-weeks old, Specific Pathogen Free (SPF), C57BL/6J male mice were purchased from The Jackson Laboratory. After one week of acclimation, mice were switched to either a western diet (WD) D12451 containing 45% lard and 20% sucrose or a matched normal diet D12450K (ND) produced by Research Diets (New Brunswick, NJ USA). Mice were maintained on these diets for 8 weeks. All animals were housed in the Laboratory Animal Resources Center (LARC) at Oregon State University in a controlled environment (12h daylight cycle) with ad libitum access to food and water. All animal experiments were approved by the Institutional Animal Care and Use Committee (IACUC) at Oregon State University and performed in accordance with the relevant guidelines and regulations as outlined in the Guide for the Care and Use of Laboratory Animals.

### Measurement of plasma and metabolic parameters

Measurements of fasting LDL, cholesterol, glucose, insulin, and leptin, glucose tolerance, liver TAGs, and weights of brown adipose tissue (BAT) and mesenteric fat pads were performed as described previously[16, 19].

### Quantification of fecal bile acids

Total fecal samples were collected for a three-day period to calculate an average fecal output for that period. Fecal material was lyophilized and ground into powder for extraction. Powdered feces (50 mg) were mixed with a 10 μl volume of bile acid internal standards (CA-d4, 10 μg/mL; DCA-d4, 10 μg/mL; GCDCA-d4, 2 μg/mL; CDCA-d4, 2 μg/mL; TCDCA-d4, 2 μg/mL). Methanol (1.5 mL) was added and samples were shaken in a Precellys homogenizer for 1 h (Bertin Instruments, Rockville, MD, USA) containing ceramic beads, centrifuged at 16,000 rpm in a microfuge for 10 min and supernatant collected. Samples were extracted two more times as described above. Supernatants were pooled, evaporated under a vacuum and reconstituted in 0.5 mL 50% methanol and randomized for data acquisition. UPLC was performed using a 1.7 μm particle, 2.1 mm × 100, CSH C18 column (Waters, Milford, MA, United States) coupled to a hybrid triple quadrupole linear ion trap mass spectrometer (AB SCIEX, 4000 QTRAP). LC and MS conditions are described previously[20]. BAs were identified by matching their retention time, isotopic pattern, exact mass of the [M-H]-ion and fragmentation pattern with a panel of authentic standards (IROA Technologies, Sea Girt, NJ, United States).

### Cell culture

Cell culture of THP-1 and IMM cells was performed as described previously [21].

### Tissue RNA extraction and library preparation

Fresh dissected tissues were flash frozen in liquid nitrogen and stored at −80°C. Libraries for Illumina sequencing were prepared as described previously [16]. Approximately 6.6 million reads were obtained per liver sample. Reads for adipose tissue and ileum, sample sequence alignment and gene counts and identification of differentially expressed genes were performed as described previously [16].

### RNA-Seq sequence alignment and gene counts

Sequences were processed to remove the adapter, polyA and low-quality bases using the fastx_clipper command in the FASTX-Toolkit v0.0.13 (FASTX-Toolkit: FASTQ/a short-reads pre-processing tools. http://hannonlab.cshl.edu/fastx_toolkit/), followed by BBDuk (bbduk.sh) parameters of k=13, ktrim=r, forcetrimleft=11, useshortkmers=t, mink=5, qtrim=r, trimq=10, minlength=20. Reads were aligned to mouse genome and transcriptome (GENCODE GRCm38) using STAR (v020201) with default parameters. Number of reads per million for mouse genes were counted using HTSeq (v 0.6.0) and quantile normalized. Differentially expressed genes were found using BRB-ArrayTools (https://brb.nci.nih.gov/BRBArrayTools/). Following sequencing, two samples from the LFD, HFD, and TXN groups, along with three samples from the XN group, were removed due to sequencing quality issues. Additionally, one mouse in the TXN group died during the study and thus was not sequenced.

### DNA extraction and 16S rRNA gene libraries preparation

Fresh fecal pellets were collected from each mouse, frozen in liquid nitrogen and stored at −80°C at the end of the feeding study. DNA was extracted as described previously[17, 18]. Sequencing of the 16s rRNA genes was performed by the Genomics and Cell Characterization Core Facility at the University of Oregon. Briefly, custom-designed PCR primers that contain dual-indexed adapters were used to amplify the V3-V4 (806R/319F) region using NEBNext® Q5® Hot Start HiFi PCR Master Mix (New England Biolabs, Beverly, MA, USA). Samples were purified by two repeated 0.8X-ratio magnetic bead clean ups to remove all traces of primer. The optimal number of cycles were determined by initial testing on a subset of samples. A no-template control and a ZymoBIOMICS D6305 microbial community DNA standard were included (Zymo Research, Irvine, CA USA). Paired-end 300 bp sequencing was performed on an Illumina MiSeq PE300 (v3 mode) with PhiX added in to about 25%. Approximately 50-60K reads per sample were achieved. Identification of gut microbial amplicon sequence variants (ASVs) followed by chimera removal was performed with DADA2 (v1.16)[22]. Taxonomy assignment was performed using the Ribosomal Database Project’s Training Set 16 and the 10.28 release of the RDP database[23]. Raw sequencing data was deposited in the NCBI Gene Expression Omnibus.

### Transkingdom multi-organ network construction and interrogation

The transkingdom network was reconstructed generally following previously described guidelines [24]. More specifically, the Mann-Whitney p-value and Fisher’s combined p-value were calculated for each measured parameter (between HFD and LFD, XN and HFD, and TXN and HFD). FDR was then calculated separately for each data type and for each tissue. P-value and FDR thresholds were then applied.

Correlations were performed within each data set (LFD, HFD, HFD+XN, and HFD+TXN). Spearman correlations were performed on the remaining parameters, followed by combining the resulting p-values using Fisher’s z transformation of correlations in the R package meta v5.5.0 [25]. Correlations were then filtered for a maximum p-value of 0.5 and combined p-value of 0.05, except in bile acids-adipose tissue, bile acids-liver, and microbiota-ileum, where a combined p-value threshold of 0.1 was used. Correlations and subsequent FDR calculations within and between each pair of data types (micro-micro, micro-gene, gene-phenotype, gene-gene, etc.) were performed separately and then integrated into a final network.

FDR was further reduced following the removal of unexpected correlations, which are caused by the violation of correlation inequalities [26]. Within-omic correlations had more stringent statistical thresholds than between-omic correlations.

We also accounted for two other network parameters. First, network sparsity which we compared the edge:node ratio of the reconstructed network to the edge:node ratio of a complete network with the same number of nodes, using the formula below.

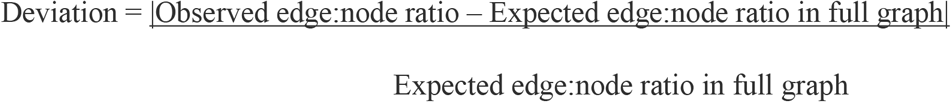

Second, to reduce the bias of positive correlations [27], we chose our statistical thresholds (below arbitrary chosen FDR, see above) that the observed negative to positive signs of correlation ratio in a network would be the closest to a full graph with no noise that fulfills correlation inequalities criteria [26].

To establish the most critical microbes for the transition to a diseased state following prolonged consumption of a high fat diet, we employed the use of bipartite betweenness centrality (BiBC). As described previously [24], BiBC is a measure of how vital a node is for information to get from one group in a network to a second group. It identifies the nodes that are the bottlenecks between groups, allowing for inference of these nodes as probably targets for treatment. This inference was then experimentally validated [21, 28]. In this study, we used the microbiota and host phenotypes as our two node groups, allowing us to identify which microbiota are the top mediators of this disease.

### Random network analysis

Random networks were created and analyzed as described previously [29]. In brief, 10,000 Erdos-Renyi random networks were created, using the same nodes in the real microbiota-adipose tissue-phenotype networks and the same number of edges (7271). BiBC was calculated between microbiota and phenotypes for each random network and the maximum node BiBC was extracted. These were then scaled on a 0 to 1 scale and compared to the actual calculated BiBC of *Oscillibacter sp*. in the reconstructed network.

### Isolation of stromal cells from visceral adipose tissue

Epididymal adipose tissue from four C57BL/6J male mice were isolated separately, mechanically dissected, then minced in Hanks’ Balanced Salt Solution (HBSS; ThermoFisher Scientific, Waltham, MA USA) containing calcium, magnesium and 0.5% BSA and 3 mg/ml collagenase type 1 (ROCKLAND, Philadelphia, PA USA). After incubation at 37°C for 1 h, the cell suspension was filtered through 100 *μ*M strainer and then washed with 20 mL DPBS containing 1 mM EDTA and 0.5% BSA (wash buffer). After centrifugation at 700 g for 15 min, the supernatant was removed, and the cell pellet was re-suspended in 5 mL of RBC lysis buffer (TONBO™ A Cytek® Brand, San Diego, CA USA) and incubated for 10 min at room temperature. Wash buffer (10 mL) was added, and the cell suspension was centrifuged again. The cell pellet was re-suspended in 1 mL FBS containing 10% DMSO and stored at liquid nitrogen until use.

### Single cell RNA seq from adipose tissue stromal cells

Stromal cells from epididymal fat tissue of four C57BL/6J male mice were isolated as described previously[21] and pooled prior to scRNA-seq. All cells were resuspended in DPBS with 0.04% BSA, and immediately processed for scRNA-seq as follows. Approximately 10,000 cells were loaded for capture onto the Chromium System using the V2 single cell reagent kit according to the manufacturer’s protocol (10× Genomics, Pleasanton, CA USA). Following capture and lysis, cDNA was synthesized and amplified as per manufacturer’s protocol (10x Genomics). The amplified cDNA from each channel of the Chromium System was used to construct an Illumina sequencing library and was sequenced on a HiSeq 4000 sequencing. Illumina base call files (*.bcl) were converted to FASTQs using CellRanger v1.3, which uses bcl2fastq v2.17.1.14. Mouse reference genome mm10 was used to align FASTQ files and transcriptome using default parameters with the CellRanger v1.3 software pipeline as previously reported; this demultiplexes the samples and generates a gene versus cell expression matrix based on the barcodes and assigns UMIs (unique molecular identifier) that enables determination of each of the individual cell from the pooled adipose tissue stromal cell samples which the RNA molecule originated.

### Single cell RNA seq data analysis

The raw gene expression matrix (UMI counts per gene per cell) was filtered, normalized, and clustered using a standard Seurat version 3.1.0 in in R (https://www.R-project.org/)[30]. Cell and gene filtering were performed as follows: Cells with a very small library size (<2500) and a very high (>0.5) mitochondrial genome transcript ratio were removed during quality check. Genes detected (UMI count > 0) in less than three cells were removed. Log normalization, clustering is performed using standard Seurat package procedures. Principal component analysis was used to reduce the number of dimensions representing each cell. The number of components from this analysis used for the elbow of a scree plot.

Based on differential expression between neighboring clusters in the samples and biological relevance, the numbers of clusters were selected. The different clusters in a sample were visualized using t-distributed Stochastic Neighbor Embedding of the principal components as implemented in Seurat. The specific cell-type identities for each cluster were determined manually using a compiled panel of available known immune cells and other cell specific marker expression.

### Additional Datasets

Obese adipose tissue single cell data [31] (GSE117176 from GSM3272967 obese ATM) was reanalyzed similarly as mentioned above to infer additional cell type information for the genes in the tissue specific network genes.

### Cell type assignments to network genes from single cell data

Adipose tissue genes from the transkingdom network were classified into specific cell types based on primarily by significant fold change in a specific cell cluster and additionally by individual genes ranked average expression in a specific cell cluster. Rules for which cell, a gene belongs to is determined by a) clusters of given cells over all other cells with -p value (<0.05) and fold change (log2FC>0.25). This was done for all genes in the adipose tissue network. It was followed as a basic rule to assign a gene to a cell type. Alternatively, a higher expression in the cell cluster (and an optional fold change (log2FC >0.25, p value <0.05) for a gene was assigned with that specific cell type. Here ranking by average expression for each gene in the clusters helps to determine its cluster specificity by higher expression in that cell type than another. This was implemented for evaluating highest cluster average expression of a gene, among all other cell clusters in network. A similar method was followed for the liver tissue network as well as to assign the cell type information to individual genes.

### Microbiota dependent gene expression in the adipose tissue

Methods were followed as detailed previously [21].

### Defining mitochondrial genes in the adipose tissue

Methods were followed as detailed previously [21]. Gene Ontology (GO) analysis was conducted using Metascape (https://metascape.org/gp/index.html#/main/step1) [32]. Adipose tissue genes in the network with TXN treatment (effect p value <0.05) were analyzed with GO biological process against mouse databases.

### Statistical analysis

Both transcriptome and microbiome data were relativized per million and quantile normalized prior to analysis. Bile acids and phenotypic data were median normalized across all samples. As the data did not follow a normal distribution, R (version and package) was used to perform Mann-Whitney comparisons between the groups (HFD vs LFD, XN vs HFD, and TXN vs HFD). Two-tailed tests were performed unless otherwise noted. Correlations were performed as described in the previous section, “Transkingdom multi-organ network construction and interrogation”. Graphpad v9.4.1 was used to perform chi-square tests. Analysis of the comparison of the observed *Oscillibacter sp*.’s BiBC compared to the distribution of BiBCs found in the random networks was performed using a one-sample Wilcoxon test with the function wilcox.test() in the R stats package v3.6.2. Additional details of statistical analyses are described in the corresponding figure legends.

## Results

### TXN more effectively restores metabolic alterations induced by HFD than XN

To determine the effects of TXN in restoring metabolic alterations induced by a HFD, we used a diet-induced mouse model of obesity and metabolic disease. We randomly assigned 10 mice to one of five groups: LFD, HFD, HFD with added XN (0.035%, low XN), HFD with added XN (0.07%, high XN) and HFD with added TXN (0.035%) for 16 weeks total [16]. We omitted the low XN dose group for downstream analyses in this study as it showed little effect on phenotypic outcomes and lost one mouse in the TXN group (n=9). In each group, we measured changes in the transcriptome of liver, ileum, and epididymal adipose tissue (adipose tissue), microbiome composition in stool, bile acid concentrations in stool, and phenotypic parameters associated with metabolic disease (**Figure 1A**). We measured weight, plasma lipid levels, plasma leptin/insulin, and plasma glucose levels as part of these phenotypic characteristics [16].

**Figure 1.**
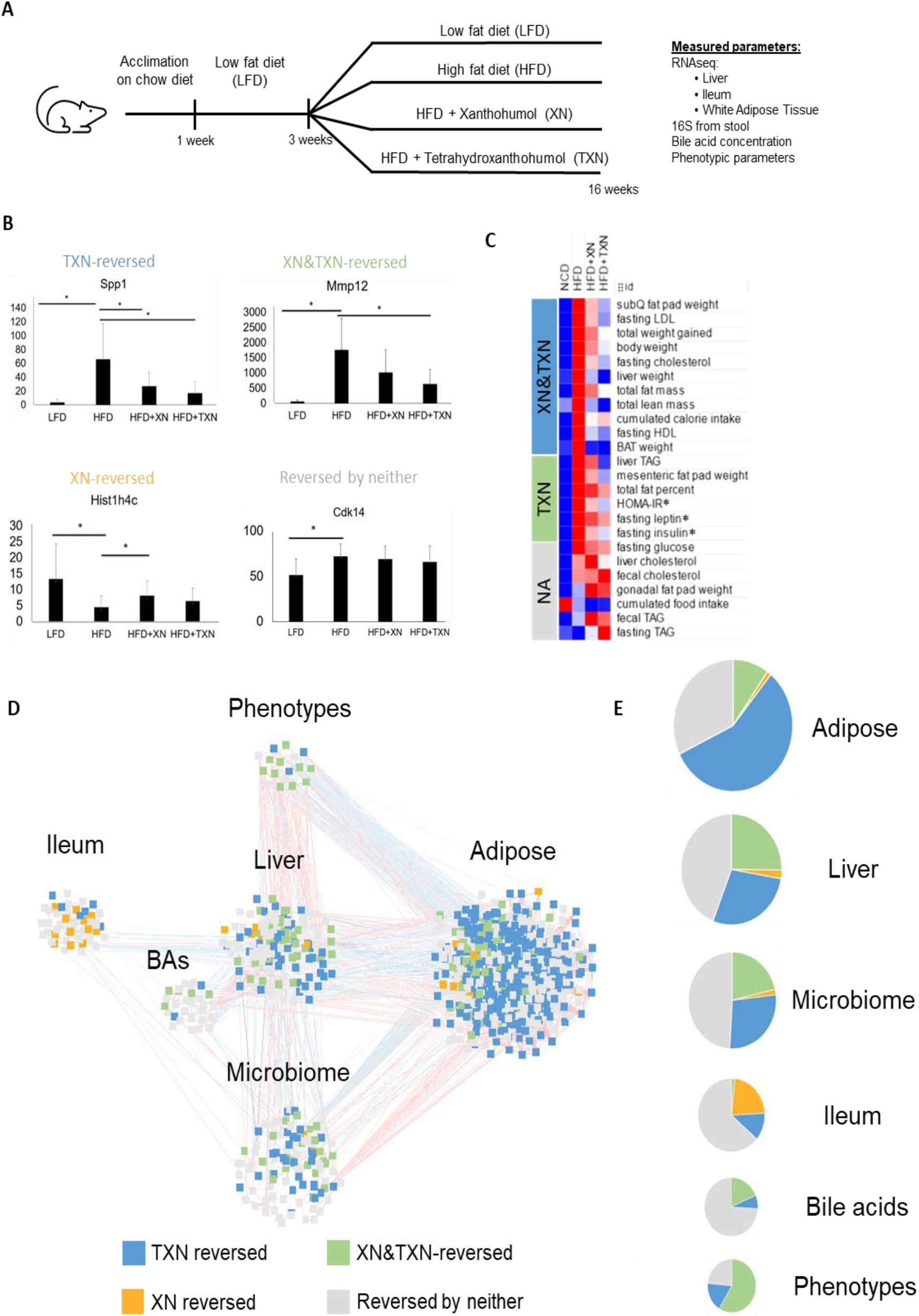
TXN primarily effects adipose tissue. A. Overview of the mouse experiments, duration, and measurements. The study contained three treatment groups: low fat control diet (LFD), high fat diet (HFD), high fat diet with XN supplementation (HFD+XN) and high fat diet with TXN supplementation (HFD+TXN). Phenotypic parameters, fecal bile acid composition, fecal 16S rRNA gene sequences, and tissue transcriptomes were determined. B. Each parameter was categorized into one of four different groups (XN-reversed, TXN-reversed, XN & TXN-reversed, reversed by neither). Shown here is an example of the expression of one gene from each category. C. The majority of measured phenotypes (especially those related to glucose homeostasis and adiposity) are restored by TXN (TXN-dependent) when compared to the direction of change in the HFD group (Mann-Whitney *P* < 0.05, * = Mann-Whitney *P* < 0.05 in HFD and < 0.1 in TXN). D. A transkingdom multi-organ network was reconstructed. Nodes in the network represent parameters that are significantly changed between HFD and LFD. Edges represent Spearman correlations between parameters (red is a positive association, blue is negative). Nodes are colored based on the treatment effect category as described in the Figure 1B examples. E. The proportion of parameters in each treatment effect category described in Figure 1D.

To elucidate differences between XN and TXN treatments, we identified those parameters reversed by either XN or TXN treatment when compared to the HFD. For example, a phenotype (e.g. body weight) increased by HFD as compared to LFD and significantly decreased by one of the treatments was considered reversed. We used this to classify all parameters into four primary categories: XN-reversed, TXN-reversed, XN and TXN-reversed, and reversed by neither (**Figure 1B**). While both XN and TXN improved 11 phenotypic parameters, including fasting LDL, fasting cholesterol, and brown adipose tissue (BAT) weight, only TXN demonstrated a significant reversal of parameters associated with diabetes, including HOMA-IR, fasting insulin and MAFLD markers like liver TAGs, mesenteric fat pad weight, and total fat percent. TXN also reversed levels of fasting leptin (**Figure 1C**). We also classified changes in transcriptomes, bile acids and microbiota into these four categories. Overall, these results confirmed our earlier observations [13, 16] that TXN is more efficacious than XN in restoring metabolic alterations caused by an obesogenic diet.

### Modeling metabolic disease via a multi-organ transkingdom network

To model HFD-induced metabolic disease, we reconstructed a multi-organ transkingdom network following our previously established method[33] (**Figure 1D**). We have previously used transkingdom networks to identify key regulatory parameters for the progression and treatment of diseases [34, 35]. Following reconstruction of the network, 17 out of 24 phenotypic parameters were represented in the network. Among the tissue transcriptomes, adipose tissue was the most prominently changed by HFD, with 1061 genes represented in the network. Liver and ileum had significantly fewer represented genes (159 and 58, respectively) (**Supplementary Table 1**). In addition, the network included 27 bile acids and 108 microbial ASVs. Strikingly, TXN alone reversed more than 50% of the genes in the adipose tissue network, while both XN and TXN reversed approximately 10% of genes (**Figure 1E**). XN alone only reversed approximately 1% of genes in the adipose tissue. We observed a similar trend of TXN reversing the expression of more genes than XN in liver as well. Compared to XN, TXN predominantly reversed the composition of the microbiome (Figure 1D & E). Both the systemic metabolic markers and prominent effects of TXN on the adipose tissue at a molecular level indicate the key mechanisms of TXN’s metabolic benefits act through the adipose tissue.

### TXN attenuates inflammatory processes involving macrophages in adipose tissue

Because 83% of the genes in the network are from adipose tissue and TXN reversed the majority (approximately 67%) of adipose tissue genes (**Figure 1E, 2A**), we determined which biological functions TXN treatment restored. Genes enriched for inflammatory response, including myeloid leukocyte activation, cytokine production, and phagocytosis were predominantly downregulated by TXN treatment (**Figure 2B**). Conversely, TXN upregulated genes enriched for metabolic processes through the restoration of lipid, amino acid and other metabolic processes (**Figure 2C**).

**Figure 2.**
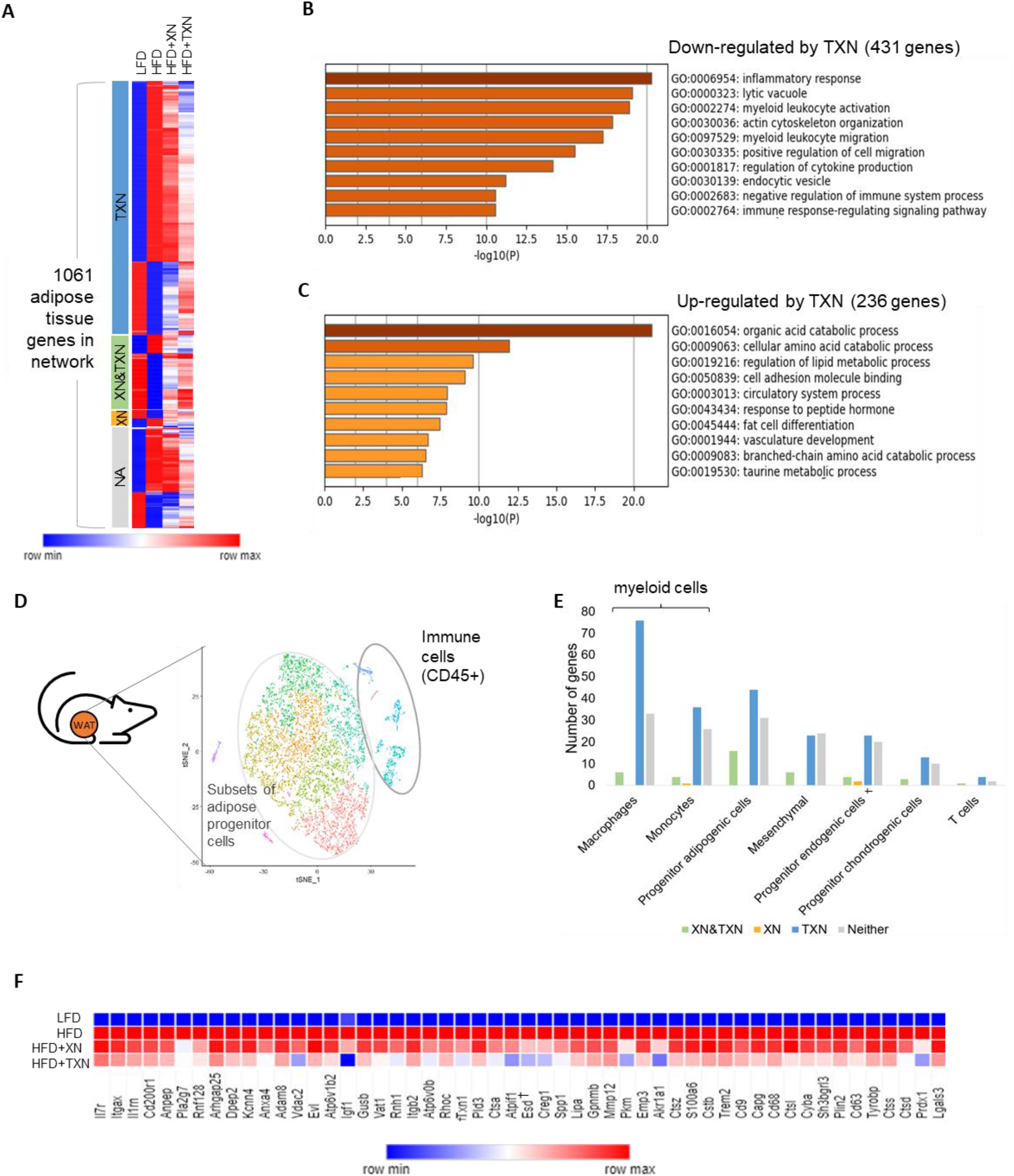
Inflammatory processes in adipose tissue are the primary processes affected by TXN. A. Heatmap representing expression of each gene in the adipose tissue subnetwork across all treatment groups. B. Metascape functional enrichment of genes down-regulated by TXN are enriched for inflammatory processes. C. Genes up-regulated by TXN are enriched for metabolic pathways. D. The t-SNE plot shows adipose tissue from high fat high sugar diet fed mice (n=5 mice, with 9,383 pooled number of cells after preprocessing for quality [Supplementary Table 5]). E. The number of genes belonging to each cell-type assignment in the adipose tissue subnetwork. TXN reverses gene expression primarily in the myeloid cell population. F. Adipose tissue network genes reversed by TXN treatment in the IR-ATM gene signature are shown. † indicates non-significance.

To investigate the reduction in inflammation, we identified those cells primarily responsible for these transcriptomic changes in the adipose tissue. To infer cell type information, we compared our whole tissue transcriptomic data with single cell RNA sequencing data from adipose tissue of mice fed an obesogenic diet [21]. Adipose tissue genes from the network were classified into specific cell types based on their average expression and fold change in specific cell clusters (**Figure 2D, Supplementary Table 5**). TXN primarily affected expression of genes assigned to myeloid cell types (macrophages and monocytes) (**Figure 2E**). These genes were enriched for inflammatory processes, including endocytosis and myeloid leukocyte migration (**Figure S1A**), and regulation of the insulin receptor-signaling pathway.

In addition to the above assignment, to confirm the cell type information, we also took advantage of a previously published dataset (GSE117176) [36] that contains single cell sequencing of mice on an obesogenic diet (**Figure S1B**). In accordance with our previous result (**Figure 2D & E**), we again found a majority of adipose tissue genes were assigned to metabolic macrophage cells (MC1) (Figure S**1C**, **left panel**). Finally, we determined if TXN altered the expression of the genes comprising the transcriptomic meta-signature of macrophages associated with metabolic disease and insulin resistance (named Insulin <u>R</u>esistance associated <u>A</u>dipose <u>T</u>issue <u>M</u>acrophages: IR-ATMs) (**Figure S1C, right panel**) [21].

Remarkably, 52 of 62 IR-ATM genes detected in the adipose tissue subnetwork were reversed by TXN treatment (**Figure 2F, Figure S1D**), supporting a role for TXN in improving the metabolic phenotype of obese mice by decreasing macrophage related inflammation in adipose tissue.

### TXN reverses microbiota-dependent IR-ATM genes

We previously showed XN requires microbiota to mediate its beneficial effects on host metabolism [18]. Therefore, we hypothesized the phenotypic changes seen in TXN-treated mice could result from communication between the gut microbiome and myeloid cells in the adipose tissue. TXN treatment increased the relative abundances of the families Verrucomicrobiaceae and Bacteroidaceae when compared to the HFD group while decreasing S24-7 (**Figure 3A**). Both beta diversity and alpha diversity were also significantly changed (**Figure 3B, Figure S2A**).

**Figure 3.**
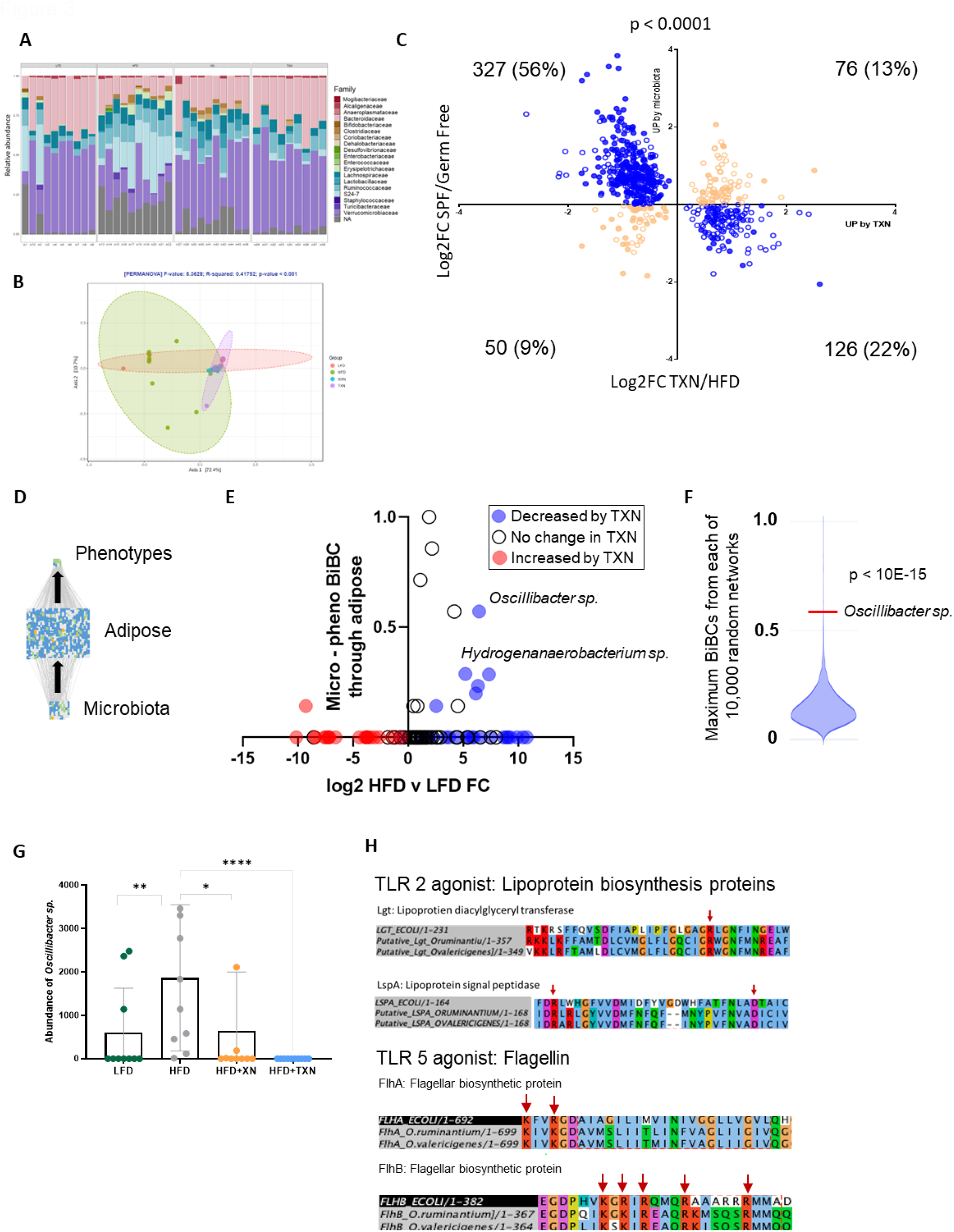
Identification of regulatory microbes causing the inflammatory response. A. Family-wise relative abundance plots of microbes across the different treatment groups. B. PCoA plot of microbiota based on treatment group. C. The microbiota dependent up- and down-regulated genes present in adipose tissue and reversed by TXN treatment. The x-axis represents log_2_ fold change TXN/HFD and y-axis represents log_2_ fold change SPF/Germ Free mice. Blue circles indicate genes whose expression was reversed by TXN treatment and microbiota-dependent. Orange circles indicate no reversal in expression direction. The filled circles were assigned to myeloid cells (two-sided Fisher’s exact test, *P* < 0.0001). D. The subnetwork derived from the original network to identify regulatory microbes that affect the host phenotypes by acting through adipose tissue. E. BiBC between microbes and phenotypes indicates *Oscillibacter sp*. and *Hydrogenanaerobacterium sp*. as two regulatory microbes whose levels are both reduced by TXN treatment. F. An analysis of 10,000 random networks with the same number of nodes and edges as the observed network revealed that the observed BiBC for *Oscillibacter sp*. was much higher than due to chance (*P* < 10E-15, one-sample t-test). G. The abundance of ASV_61 (identified as *Oscillibacter* sp.) in each treatment group. Two-tailed Mann-Whitney P values are shown; brackets indicate the results for specific comparisons (\**P* ≤ 0.05; \*\*\*\**P* ≤ 0.0001). H. ClustalO multiple sequence alignment shows TLR2/TLR5 agonist protein sequences aligned with respective *E. coli* sequences containing conserved domains, which also have same functional residues. The red arrows indicate the functional residues in the protein sequence alignment.

To identify how TXN acts on host functionally via the microbiota, we used a previous method for discriminating between microbiota-dependent and -independent effects of antibiotics [37]. For this, we established which TXN transcriptomic alterations in adipose tissue are dependent on microbiota using gene expression data from germ free and SPF mice. We found a large overlap between microbiota-dependent genes and those affected by TXN treatment (**Supplementary Table 3**). Specifically, out of 667 genes reversed by TXN in adipose tissue, 78% were also controlled by microbiota, but in the opposing direction (**Figure 3C**). Of note, a majority of these genes were myeloid-specific (**Figure 3C, Supplementary Table 2**). As expected, the signature genes for IR-ATMs overlapped with microbiota-dependent genes and 41 of them showed a significant reversal by TXN treatment. Taken together, these findings indicate TXN reverses the effect of HFD-associated microbiota on adipose tissue gene expression.

### *Oscillibacter* sp. is one of the key microbes mediating the beneficial effect of TXN

To identify microbes responsible for the upregulation of the aforementioned IR-ATM signature genes in HFD-fed mice, we restricted our transkingdom network analysis to only the microbiota, adipose tissue transcriptome and metabolic phenotypes. Next, we inferred key microbes with causal contribution to systemic metabolic phenotypes using bipartite betweenness centrality (BiBC). Briefly, BiBC measures the “bottleneck-ness” of a node between two node groups [24]. A high BiBC value indicates a node with regulatory potential that functions as a causal agent in the model under study. In this study, BiBC was calculated between phenotypic parameters and the microbiome through the adipose tissue (**Figure 3D**). Microbes of the *Oscillibacter* (ASV_61) and *Hydrogenanaerobacterium* (ASV_70) genera appeared as the top potential causal microbes, as indicated by their decrease with TXN treatment, relatively high BiBC, and high fold-change in HFD (**Figure 3E & 3G, Figure S2B**). Comparing these results with random networks, we found *Oscillibacter sp*. is both the top ranked “bottleneck” (BiBC) node and also a highly statistically significant observation (one sample Wilcoxon test p < 10^−15^). (**Figure 3E & 3F)**. After aligning the 16S sequence of ASV_61 to NCBI’s RefSeq database using BLAST, we identified *O. ruminantium* (OR) and *O. valericigenes* (OV) as top hits, respectively. These species are >94% similar, compared to the next most similar taxa (<90%) (**Figure S2C**). Taken together, these findings indicate TXN’s beneficial effects on adipose tissue and systemic metabolism are mediated by reducing levels of *Oscillibacter sp* (**Figure 3G**).

In a recent study with a diet-induced T2D mouse model, we found *Oscillibacter* spp. promotes insulin resistance by increasing metabolically damaging macrophages in adipose tissue [21]. There, we demonstrated TLR2 and potentially TLR5 agonists produced by this microbe predominantly mediate the effect of OV on macrophages. In the current study, we cannot definitively pinpoint which of two species of *Oscillibacter* are present due to limitations of 16S amplicon sequencing; however, we verified both *O. ruminantium* and *O. valericigenes* are potentially capable of producing Tlr2 and Tlr5 agonists. We identified and aligned the probable *Oscillibacter* species’ protein sequences involved in the biosynthetic processes of TLR2 and TLR5 agonists to known lipoprotein synthesis proteins (*Lgt*, lipoprotein diacylglyceryl transferase (PDB accession code 5AZB) [38] and *LspA*, lipoprotein signal peptidase from *E. coli*) and flagellin synthesis with export apparatus (*FlhAB*, flagellar biosynthetic proteins), respectively (**Figure 3H**). The highly conserved domains in these alignments indicate these putative proteins are likely functional. Reducing the abundance of these microbes in the gut could reduce the level of TLR agonists, leading to diminished inflammatory responses.

As described above, we inferred that TXN decreases the IR-ATM signature by reducing *Oscillibacter*. Furthermore, in our recent study [21] we showed that IR-ATMs negatively affect systemic metabolism via damage of mitochondria in adipose tissue. Thus, we hypothesized TXN may restore adipose tissue mitochondrial function in mediating its beneficial downstream effects. To support our hypothesis, we determined the effect of TXN on mitochondrial gene expression in the adipose tissue. First, we found genes reversed by TXN in the adipose tissue were enriched for mitochondrial processes such as antioxidant activity and oxidoreductase activity (**Figure S2D**). Next, we identified 87 mitochondrial genes (MitoCarta 3.0, MitoGenes [21, 39] in the adipose tissue network that were reversed by TXN. Among the genes predicted to mediate the effect of myeloid cells on systemic metabolism (high BiBC between myeloid cells and phenotypes), we observed a trend of enrichment for mitochondrial function. For example, among these predicted regulatory genes, there were several genes with well-known mitochondrial function such as thioredoxin reductase [40], glutathione transferase [41], and many coenzyme A-related genes [42] (**Figure S2E**). Taken together, our previous findings and these results indicate TXN treatment may improve mitochondrial function in the adipose tissue, similar to what we previously reported for the liver [43].

### Macrophage genes induced by *Oscillibacter* are inhibited by TXN

To assess the inflammatory effects of OV, we exposed two mouse macrophage cell lines, RAW 264.7 and IMM (immortalized mouse macrophage cells) to supernatant from this microbe (**Figure 4A**). OV supernatant altered the transcriptome of 2,241 genes in both cell lines. Of those, TXN reversed the expression of 325 genes in RAW 264.7 cells that belonged to the adipose tissue network and were microbiota-dependent. In IMM cells, 268 genes fit the same criteria (**Figure S3A**). Overall, of the 667 adipose tissue genes reversed by TXN, 157 overlapped with the genes regulated by OV supernatant in both IMM and RAW264.7 cells (**Figure 4A**). Specifically, supernatant exposure increased IR-ATM signature genes including *Itgax, Adam8, Spp1, Plin2*, and *Prdx1* in both cell lines (**Figure 4C, Supplementary Table 3**), as well as *Atf3* (**Figure 4C, Supplementary Table 3**), a transcription factor that controls critical effector molecules of these macrophages [21]. Of note, 79 of these 157 genes, including many IR-ATM signature genes were significantly increased in both OV supernatant-exposed macrophage cell lines, while the same genes were reversed by TXN treatment *in vivo* in the adipose tissue myeloid cells **(Figure 4B, figure S3B**). The effect of OV explains around half of the microbiota-dependent transcriptomic alterations in macrophages caused by TXN *in vivo* **(Figure S3A)**.

**Figure 4.**
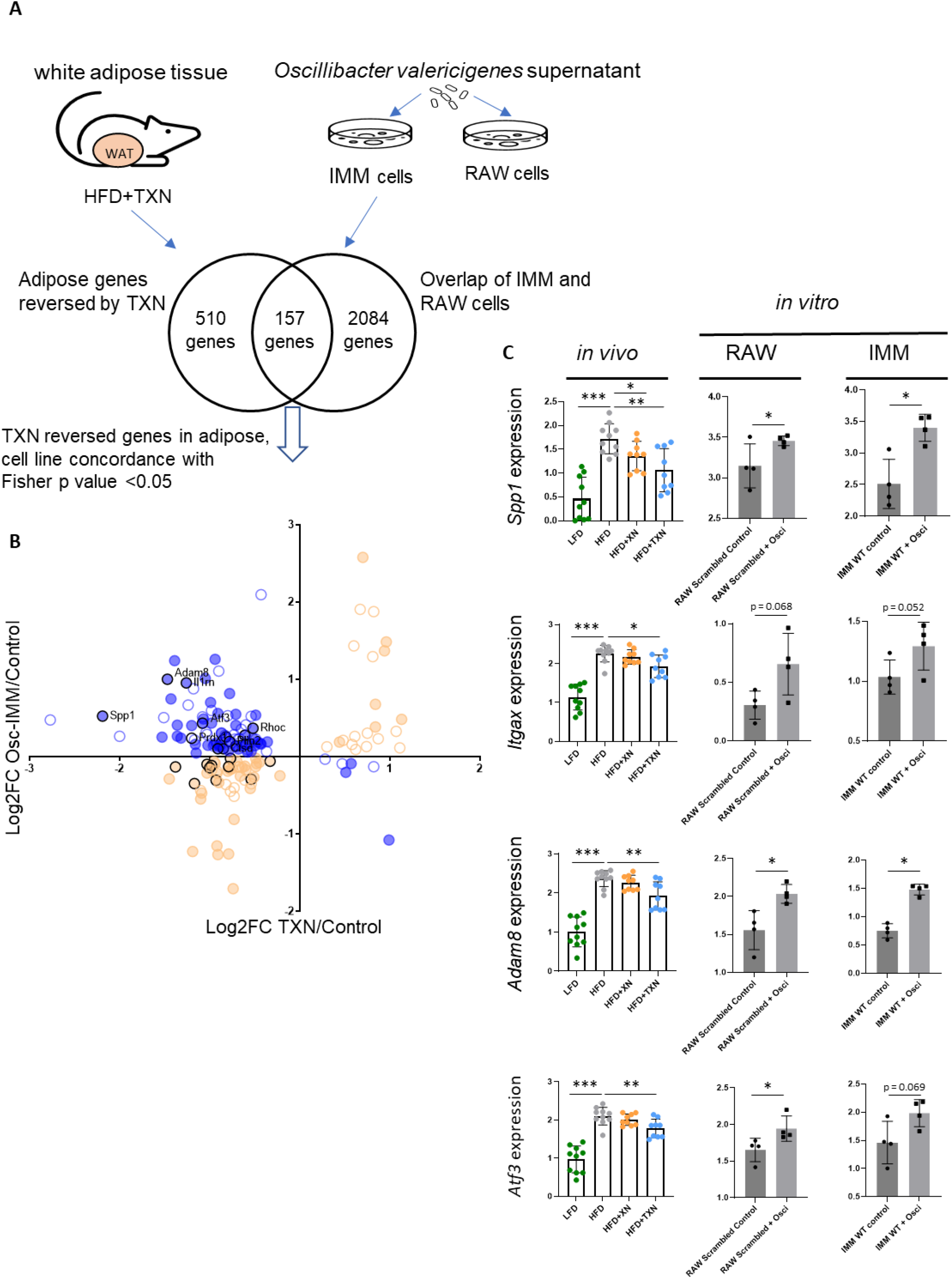
*O. valericigenes* treatment increases the expression of IR-ATM genes that are significantly reversed by TXN in macrophage cell lines. A. Treatment of macrophage cell lines IMM and RAW 264.7 with *O. valericigenes* supernatant increased the expression of several important IR-ATM genes (one-sided Fisher’s exact test; BH-adjusted *P* < 0.05) present in the overlap of the Venn diagram. The same genes are reversed by TXN treatment (Mann-Whitney test; *P* < 0.05) in the adipose tissue of mice fed an HFD (HFD+TXN). B. The XY-plot shows the genes significantly reversed by TXN treatment concordantly in both macrophage cell lines with individual comparison *P* < 0.1. Blue circles indicate those genes reversed by TXN treatment and microbiota-dependent. Orange circles indicate no change in expression direction. The filled circles indicate those genes assigned to myeloid cells. The black circles indicate IR-ATM signature genes (one-sided Chi-square test, *P* < 0.05). C. Expression of representative genes reversed (decreased) by TXN treatment in the adipose tissue but increased with *O. valericigenes* treatment of macrophage cell lines. * Mann-Whitney p-value < 0.05; ** p-value < 0.01; *** p-value < 0.001.

To support our in vitro findings, we compared expression of adipose tissue genes from mice treated with TXN versus those from mice fed a control diet and supplemented daily with OV (10^9^ CFU) for 4-5 weeks. We found that 75 (73 up- and 2 down-regulated) out of 99 genes modulated by OV treatment were all reversed by TXN treatment. Of these 75 genes, 17 (including *Mmp12, Lgals3*, and *Plin2*) belonged to the IR-ATM signature (**Figure 5A-C**). We postulate that other microbes such as *Hydrogenanaerobacterium* (**Figure 3E**) could further promote inflammatory metabolic macrophages and related phenomenon. Taken together, our results indicate the interaction between the gut microbiota and adipose tissue comprise the key cellular and molecular events mediating beneficial effects of TXN on systemic metabolism. Specifically, we posit that reduction of *Oscillibacter* (and potentially other bacterial producers of Tlr2/5 ligands) levels and the consequent decrease and/or reprogramming of metabolically harmful IR-ATMs represents one of the main mechanisms of TXN’s therapeutic action in MetS (**Figure 6**).

**Figure 5.**
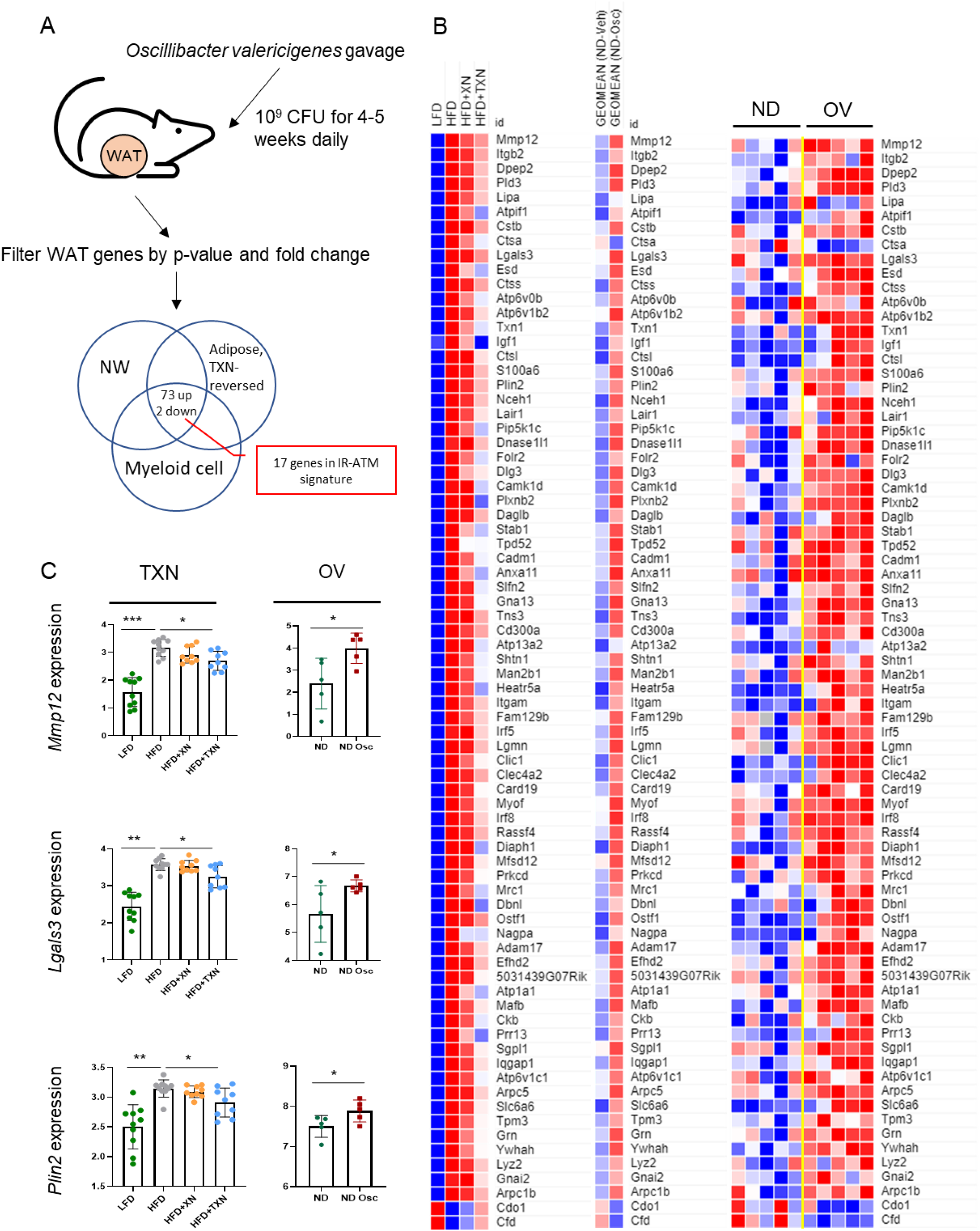
*O. valericigenes* treatment increases the expression of IR-ATM genes in mouse adipose tissue that are significantly reversed by TXN treatment. A. Experimental outline for *O. valericigenes* gavage and subsequent gene expression from the adipose tissue (n = 5 per group). B. Expression of adipose tissue genes increased by an HFD and reversed by TXN treatment in mice (left panel) vs. the same adipose tissue genes following *O. valericigenes* supplementation of mice (right panel). The middle panel shows the geometric mean of the samples in the right panel. C. Examples of adipose tissue genes (in the IR-ATM signature) decreased by TXN treatment but induced by *O. valericigenes* supplementation *in vivo*. * Mann-Whitney p-value < 0.05; ** p-value < 0.01; *** p-value < 0.001.

**Figure 6.**
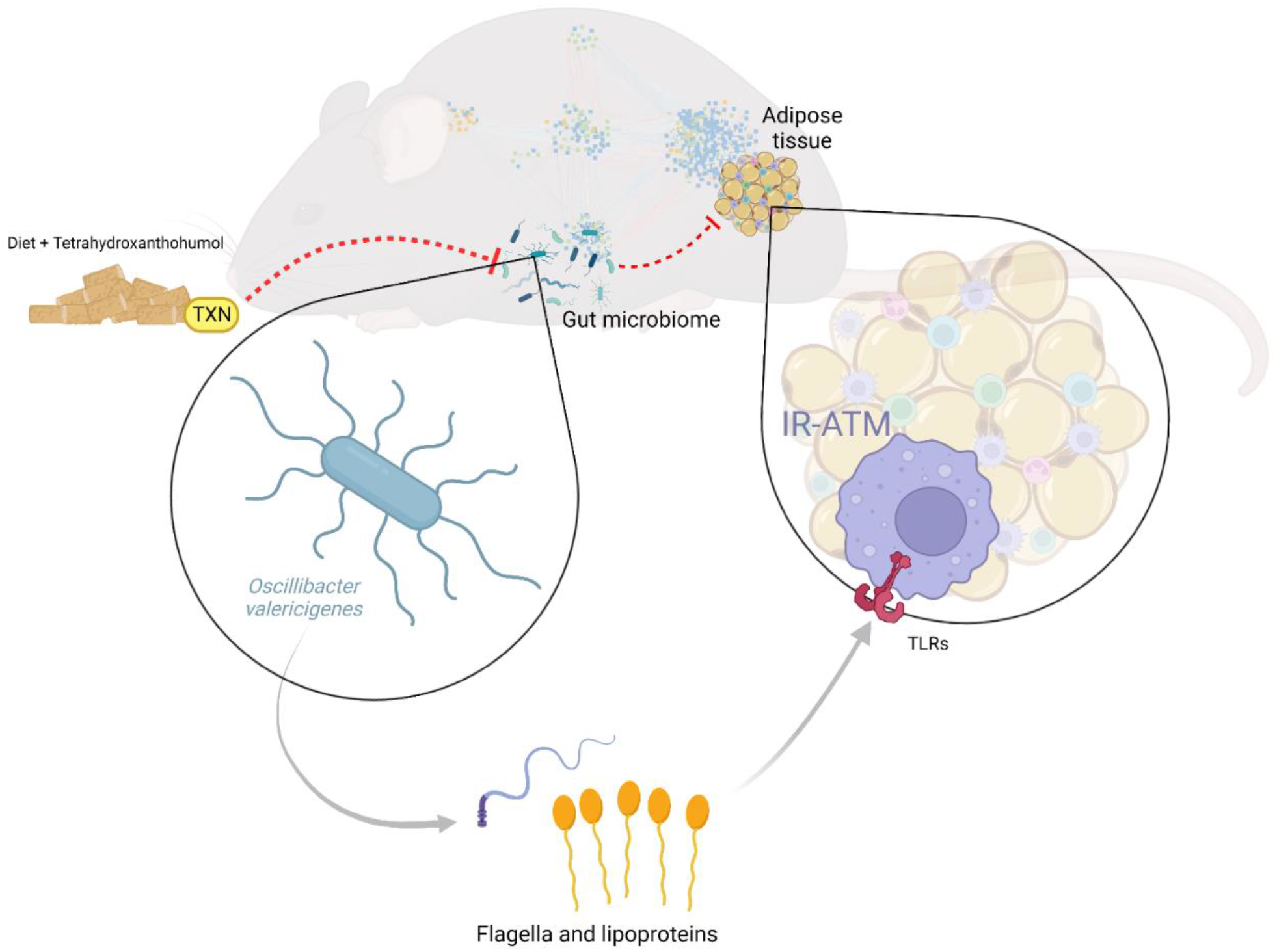
Summary Figure. Consumption of TXN effects the composition of the gut microbial community (e.g. reduces *Oscillibacter* sp.) which, in turn, decreases metabolically damaging adipose tissue macrophages that result from an obeso-/diabeto-genic diet.

## Discussion

XN and its derivatives mitigate diet-induced obesity-related characteristics of MetS in mice by improving impaired glucose and lipid metabolism [19, 43-45]. XN also exhibits hepatoprotective effects in animal models of MAFLD and other liver-damaging conditions [46, 47]. Accumulating studies indicate these prenylated flavonoids may mediate their beneficial effects via polypharmacological mechanisms of action, which could increase their effectiveness in treating disease. Previous studies link the anti-obesity and MetS benefits of XN and its derivatives to possibly increasing energy expenditure through mild mitochondrial uncoupling and increasing locomotor activity [43, 48, 49]. Xanthohumol improves diet-induced obesity and hepatosteatosis by inhibiting expression of and inactivating maturation of sterol regulatory element-binding proteins, thus reducing the expression of genes involved in fatty acid and cholesterol biosynthesis [14, 50]. Evidence also suggests xanthohumol and even more so, TXN, attenuates hepatosteatosis by acting as antagonists of PPARγ [16]. Improvement in glucose homeostasis by XN was attributed to AMPK activation in the mouse liver [13, 51]; however, we did not observe this with TXN [13]. We also found improvements in glucose metabolism by XN required the microbiota [18]. Furthermore, administration of XN and TXN altered the gut microbiota composition and bile acid metabolism in mice implicating these changes in mediating the benefits of XN and TXN [17]. In particular, administering TXN was very effective in changing gut microbiota composition, bile acid metabolism, and reducing adipose tissue inflammation [17]. Despite these numerous studies, there is no consensus which organs and molecular mechanisms contribute the most to the beneficial effects of XN and its derivative TXN.

In this study, to delineate the beneficial effects of TXN, we used a multi-omics transkingdom network analysis from in vivo data obtained from diet-induced obese murine models and validated the outcome of the analysis with additional in vitro and in vivo data. TXN acted predominantly on adipose tissue with approximately two-thirds of DEGs altered by TXN treatment. Of those genes whose expression was decreased by TXN, there was significant enrichment for inflammatory processes, implicating myeloid cells, especially macrophages, as an important factor in the disease in part dependent on microbiota.

The connection between gut microbes and obesity related inflammation leading to MetS remains a poorly characterized phenomenon [52]. In a recent publication [21], we described expansion of *O. valericigenes* among a few other gut microbes in mice fed a high fat, high sugar diet, which increased Mmp12-positive macrophages with a particular transcriptomic signature indicative of a unique cell population of IR-ATMs. In this study, using an HFD in mice in a different facility, our unbiased investigation identified inflammatory macrophages with a very similar signature in the adipose tissue.

Causal inference via our transkingdom network analysis revealed *Oscillibacter* spp. as a key microbe mediating the effects of TXN in decreasing inflammation in adipose tissue and improving symptoms of MetS. From our analysis, we inferred several microbes, including *O. valericigenes* that could activate TLRs and experimentally demonstrated this activation increased expression of numerous IR-ATM genes in vitro in RAW 264.7 and IMM cell lines and *in vivo* in adipose tissue of mice fed control diet. This activation is consistent with the molecular machinery of this pathobiont. We found *Oscillibacter* contained genes encoding proteins with high sequence identity to lipoprotein synthesis pathway proteins responsible for synthesizing lipoprotein, a TLR2 agonist [53], along with flagellin synthesis and assembly machinery proteins responsible for synthesizing flagellin, a TLR5 agonist [54]. TXN-treatment decreased expression of these proinflammatory genes, characteristic of IR-ATMs in adipose tissue of mice fed an HFD. Thus, these experiments support TXN’s mechanism of action in the adipose tissue is predominantly related to dampening macrophage-mediated inflammation. Spp1, Cd9, Trem2 and Mmp12 [21, 55-58] and other well-known IR-ATM genes were among the ones increased by the microbial supernatant while TXN significantly decreased their expression in adipose tissue. Taken together, these findings reveal that TXN benefits the host by acting on IR-ATMs via changes in the composition of the gut microbiota. These findings may have translational potential as we previously showed this macrophage signature shows a strong overlap with that found in a meta-analysis of T2D patients [21]; suggesting TXN treatment could benefit obese patients with MetS conditions.

Current literature provides compelling evidence for a model of MetS pathogenesis that involves more than a few microbes acting on different mammalian pathways and organs [59-61]. Accordingly, while the sample size of this study allowed us to infer only one top bacterium (*Oscillibacter*) clearly contributing to the beneficial effect of TXN, we also observed a similar functional trend for a few other microbes decreased by TXN (**Figure 3E**). Some (e.g. *Hydrogenanaerobacterium*) were previously reported by us and others [62] to be associated with MetS. Moreover, a previous study demonstrated that at least some of the positive outcomes of XN are accomplished by increasing *A. muciniphila*, which we also found increased by TXN [18]. These results suggest that when microbiota mediate the effect of a given treatment, the specific cellular/molecular mechanism would strongly depend on the presence/absence of causally contributing microbes in the underlying microbial community.

In conclusion, XN and its derivatives mediate their beneficial effects against obesity and MetS through several mechanisms of action. However, to the best of our knowledge, this is the first study identifying a predominant mechanism of action for TXN in reducing microbial community members (e.g. *Oscillibacter* sp.) which promote metabolically damaging adipose tissue macrophages in response to an obeso-/diabetogenic diet. Though we identified a new mechanism explaining TXN’s benefits in adipose tissue, one should note that TXN also affects intestine, liver, bile acids and additional microbiota, warranting further investigation in future studies.

## Acknowledgments

We thank Jamie Pennington, Scott Leonard, Dr. Wenbin Wu for their assistance and Drs. Amiran Dzutsev and Giorgio Trinchieri for help with scRANseq data generation. The National Institutes of Health (NIH grants 5R01AT009168 to AFG, CSM, and JFS and 1S10RR027878 to JFS; DK103761 to NS), the Linus Pauling Institute (LPI), the OSU College of Pharmacy, Hopsteiner, Inc, New York (JFS), and the OSU Foundation Buhler-Wang Research Fund (CLM) supported this research. The Marion T Tsefalas Graduate Fellowship from the LPI, the ZRT Laboratory Fund for the LPI, and the Charley Helen, Nutrition Science and Margy J Woodburn Fellowships from the School of Biological and Population Health Sciences at OSU supported YZ.

## Author Contributions

Conceptualization, Data curation, Formal analysis, Validation, Investigation, Visualization, Methodology, Writing - original draft, Project administration, Writing - review and editing (NKN, YZ, JP); Conceptualization, Resources, Data Curation, Formal analysis, Supervision, Funding acquisition, Investigation, Visualization, Methodology, Writing - original draft, Project administration, Writing - review and editing (AFG, AM, NS); Conceptualization, Resources, Funding acquisition, Methodology, Project administration, Writing - review and editing (JFS, CSM); Conceptualization, Data curation, Formal analysis, Validation, Investigation, Methodology, Project administration, Writing - review and editing (CLM, GB); Resources, Visualization, Methodology, Writing - review and editing (TJS); Data curation, Formal analysis, Validation, Investigation, Methodology, Writing - review and editing (AAM); Investigation, Writing –review and editing (CPW).

## Availability of data and materials

Raw and processed data has submitted to Gene Expression Omnibus (GEO) [63]. The sequencing of epidydimal adipose tissue in mice fed a normal diet with/without *O. valericigenes* gavage can be found under the GEO accession number GSE215226. The adipose tissue single cell data is available as a supplementary file. RNA-Seq data of IMM cells treated with Oscillibacter supernatant: GSE203488. RNA-Seq data of RAW 264.7 cells treated with Oscillibacter supernatant: GSE203516. RNA-Seq data of liver of mice: GSE164636. Microbiome 16S rRNA DNA sequencing: Sequence Read Archive SUB10676151.

**Supplementary Table 1 –** Number of parameters in each measured category. See Figure 1C for more information on how categories were assigned.

**Supplementary Table 2** – Summary table of parameters, including statistical information, cell type information, microbiota dependence/independence for genes, abundance of microbes, and network information

**Supplementary Table 3** – IMM/RAW cell with Oscillibacter supplementation (processed data: GSE203488, GSE203516)

**Supplementary Table 4** – Results of in vivo validation of *O. valericigenes* gavage

**Figure S1.**
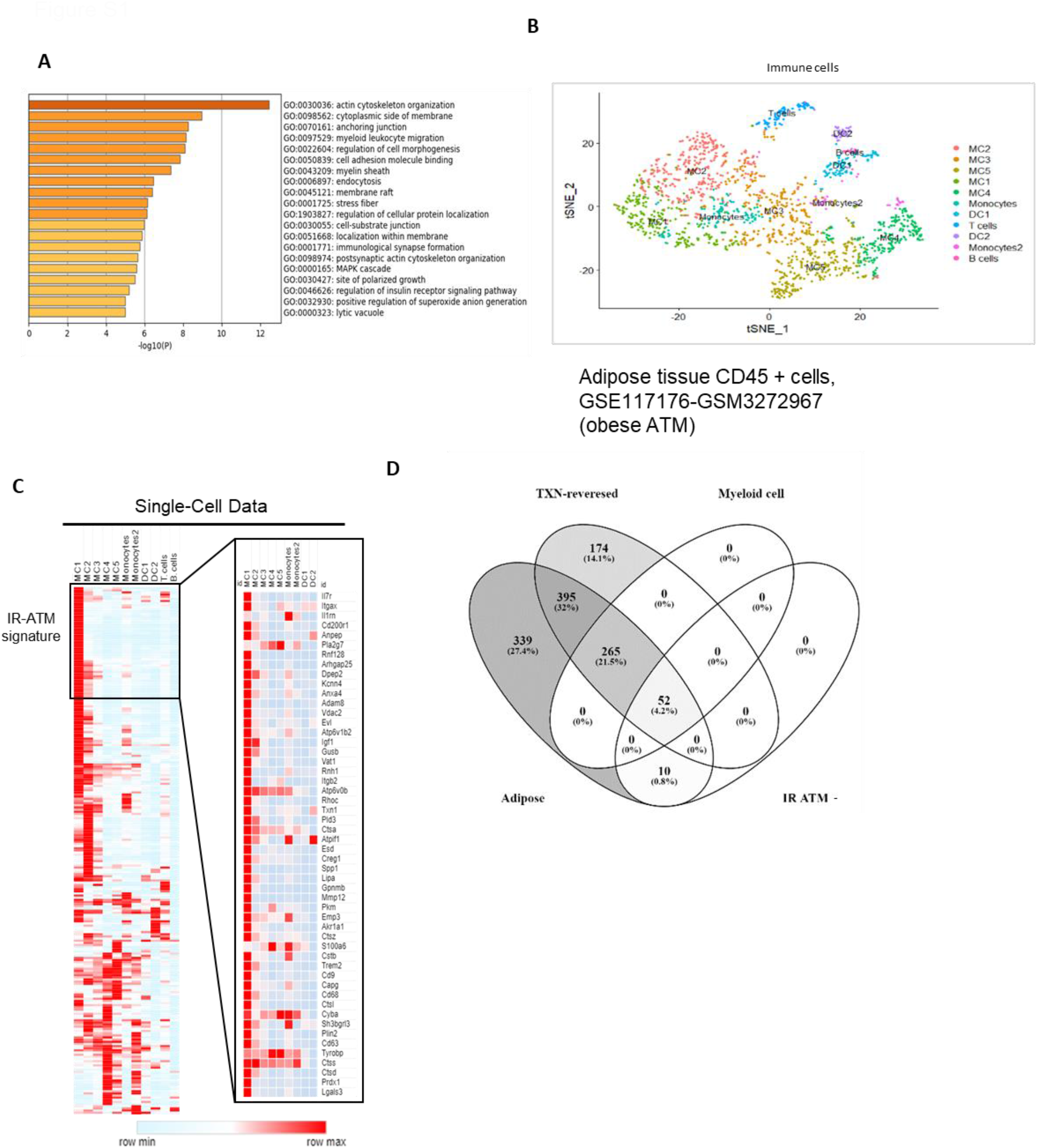
Myeloid cells are the primary cell type affected by TXN in the adipose tissue. A. Functional enrichment of genes belonging to myeloid cells and reversed by TXN treatment indicates significant enrichment of myeloid cell activation, endocytosis and insulin receptor pathways. B. T-SNE plot shows re-analyzed clusters of different cell types from a previously published dataset (GSE117176) containing single cell sequencing of murine adipose tissue stromal cells on a high fat diet. The immune cell clusters are labeled based on the specific immune cell markers derived from the reanalysis. C. Average expression from the validation single cell study (GSE117176) for each of the genes assigned to a specific cell type is shown. Many of the adipose tissue genes in our whole tissue RNA-seq study were assigned to myeloid cells, primarily the metabolic macrophage (IR-ATM) type, named here ‘MC1’ cells. The zoomed in heatmap in the right panel shows the higher average expression of IR-ATM signature genes in the MC1 group. D. The Venn diagram represents the adipose tissue genes from the network and shows the genes reversed by TXN that were assigned to myeloid cells and/or IR-ATMs. The IR-ATM gene signature is significantly reversed by TXN (52/62) as shown.

**Figure S2.**
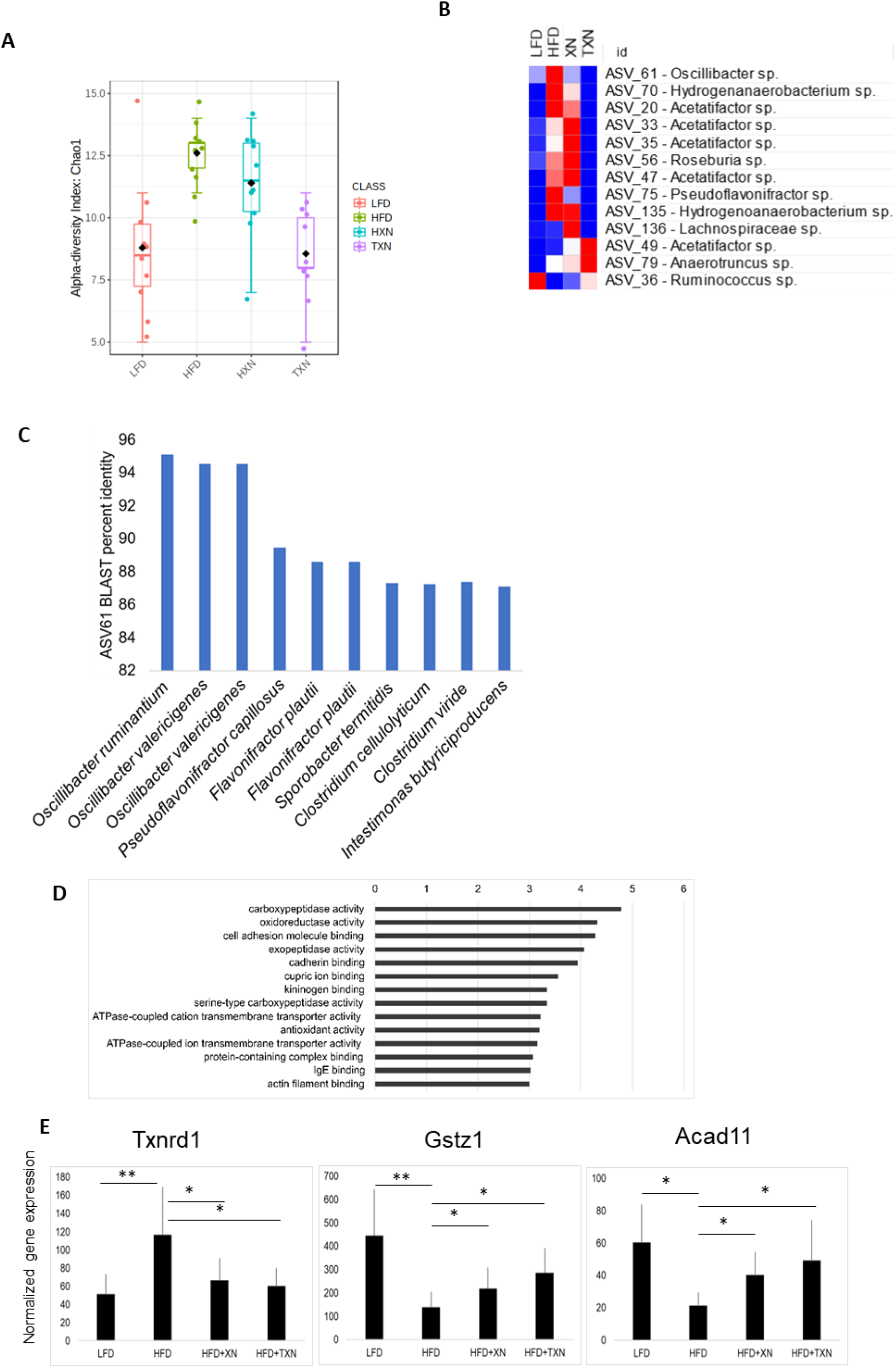
TXN treatment alters fecal microbiota composition and mitochondrial gene expression. A. Comparison of alpha diversity across treatment groups. B. Abundance of ASVs for each treatment group. ASVs plotted are from those in the top BiBC in Figure 3E. C. BLAST percent identity comparison of ASV61 indicates high similarity to both *O. ruminantium* and *O. valericigenes* as the top three hits. Subsequent hits show a much-reduced similarity. Notably, none of the subsequent hits are from the *Oscillibacter* genus. D. The top BiBC mitochondrial genes in the adipose tissue network are enriched for oxidoreductase activity and antioxidant activity. E. Expression levels for representative mitochondrial genes significantly reversed by TXN that comprise top BiBC in the adipose tissue network.

**Figure S3.**
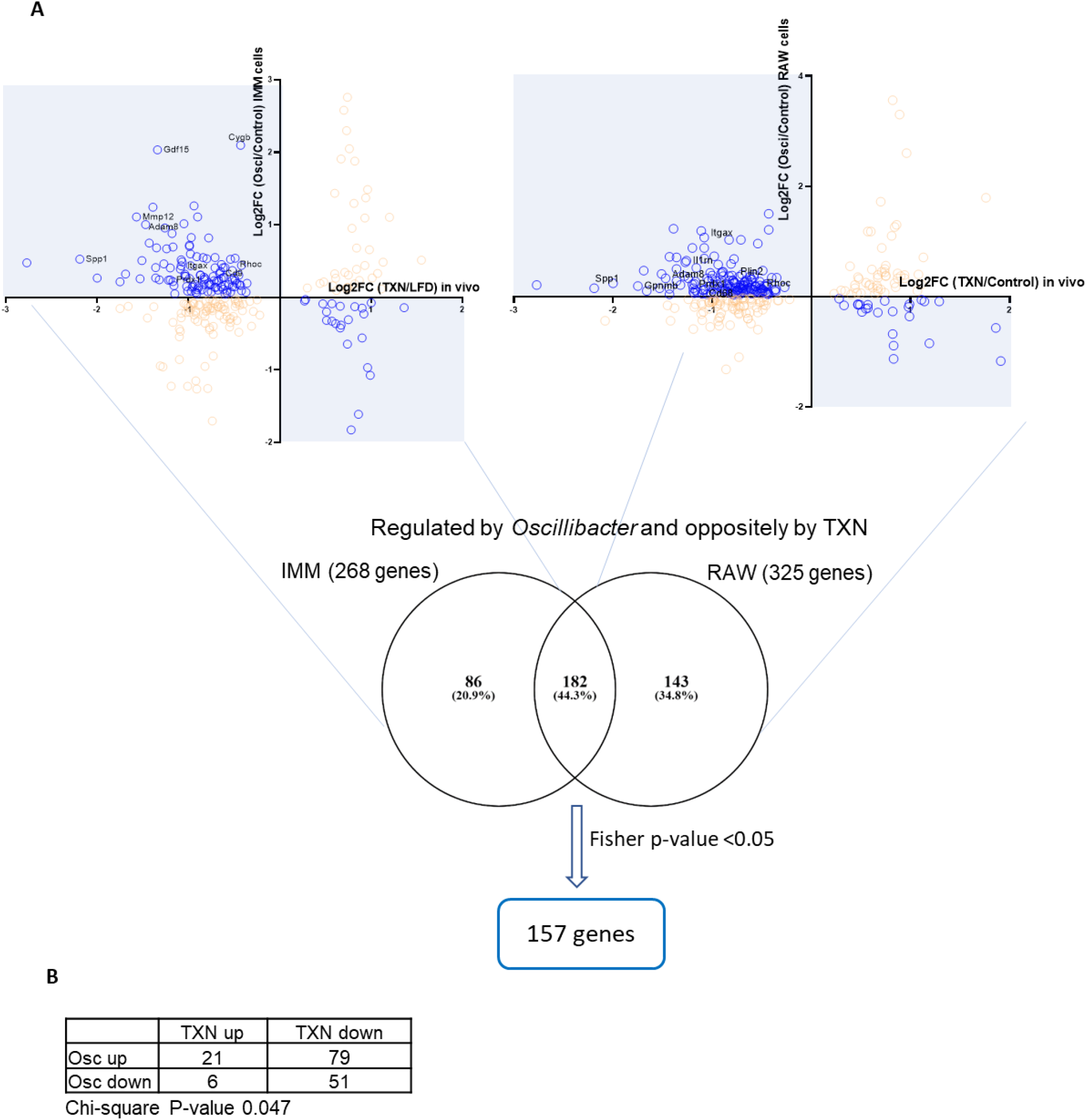
Identification of overlapping gene expression between adipose tissue from TXN-treated mice vs. *O. valericigenes* supernatant-treated macrophage cell lines. A. Finding the overlap of the genes differentially expressed in *O. valericigenes* treated IMM cells (graph on top left) and *O. valericigenes* treated RAW 364.7 cells (graph on top right) that were differentially expressed in the opposite direction with TXN treatment in vivo (blue shaded area). Circles are genes assigned to the TXN-reversed category in vivo. Mann-Whitney P < 0.1, Oscillibacter treated vs. untreated cells. Meta-analysis of the IMM and RAW 264.7 cells yields 157 significant genes (Fisher’s combined P < 0.05). B. Chi-square analysis results for Figure 4A.

## Notes

### Competing Interest Statement

The authors have declared no competing interest.

